# weIMPUTE: A User-Friendly Web-Based Genotype Imputation Platform

**DOI:** 10.1101/2023.08.10.552759

**Authors:** You Tang, Mingliang Li, Zhuo Li, Qi Li, Xiaodong Hu, Jun Yu, Jian Lin, Chunguang Bi, Helong Yu, Meng Huang

**Affiliations:** Electrical and Information Engineering College, Jilin Agricultural Science and Technology University, Jilin, Jilin, 132101, China; College of Information and Control Engineering, Jilin Institute of Chemical Technology, Jilin, Jilin, 132101, China; College of Information Technology, Jilin Agricultural University, Changchun, Jilin, 132101, China; Guangzhou College of Technology and Business School of Engineering, Guangzhou, Guangdong, 510000, China

**Keywords:** Imputation, phasing, GUI, genotype, web based

## Abstract

Genotype imputation is a critical preprocessing step in genome-wide association studies (GWAS), enhancing statistical power for detecting associated single nucleotide polymorphisms (SNPs) by increasing marker size. In response to the needs of researchers seeking user-friendly graphical tools for imputation without requiring informatics or computer expertise, we have developed weIMPUTE, a web-based imputation graphical user interface (GUI). Unlike existing genotype imputation software, weIMPUTE supports multiple imputation software, including SHAPEIT, Eagle, Minimac4, Beagle, and IMPUTE2, while encompassing the entire workflow, from quality control to data format conversion. This comprehensive platform enables both novices and experienced users to readily perform imputation tasks. For reference genotype data owners, weIMPUTE can be installed on a server or workstation, facilitating web-based imputation services without data sharing. weIMPUTE represents a versatile imputation solution for researchers across various fields, offering the flexibility to create personalized imputation servers on different operating systems.

## Introduction

The advent of high-throughput sequencing data has revolutionized research endeavors, leading to significant advances in areas such as genome-wide association studies (GWAS) and genome selection (GS). However, there remains a crucial challenge of efficiently imputing low-density datasets to high-density ones. Fully harnessing the potential of whole genome sequencing technology to increase sample sizes and enhance statistical power in GWAS, as well as reducing the cost of GS in animal and plant breeding, necessitates a user-friendly imputation pipeline that does not require extensive command line knowledge or expertise in the Linux environment. Moreover, establishing an imputation server offers an effective means to leverage computing resources for large datasets while securely providing public imputation services.

While certain free accessible services, like TOPMed ^2^, accommodate over 200,000 individuals, they may not be universally available to all researchers. Another popular option, the Michigan imputation server (MIS), caters to human genetic researchers but supports only Eagle ^3^ and Minimac4 ^1^ as imputation tools and lacks data preparation functions such as format conversion and quality control. Consequently, researchers without prior experience in data cleaning or those from non-human fields seek an alternative, easy-to-use, and integrated imputation solution to offer personalized imputation services to their communities.

Genotype imputation is a well-established and continuously evolving method. Dealing with large reference datasets often involves separating the phasing and imputing processes. Widely used software, such as Minimac4, Eagle2, Beagle5, SHAPEIT, and IMPUTE2, each possesses unique features for genotype phasing or imputation ^1,3–7^. While some tools like Eagle and Beagle can conduct phasing or imputing without a reference population, others, like SHAPEIT-IMPUTE2, are designed to handle admixture populations. Additionally, methods like Minimac4 require homogeneous populations, and except for Beagle, which is cross-platform and written in Java, the other three software can only be used in a Linux environment. To cater to diverse research needs, we have developed the weIMPUTE imputation platform. weIMPUTE offers users the flexibility to choose between two pre-phasing software options, Eagle2 or SHAPEIT, and three imputation software options, Minimac4, Beagle5, and IMPUTE2. Leveraging Docker technology, weIMPUTE enables seamless installation and usage across different operating systems. The platform also integrates quality control (QC) and post-filtering, streamlining the imputation process in a one-stop service. Additionally, weIMPUTE facilitates the parallel launch of multiple jobs, efficiently utilizing computing resources. Through comprehensive testing, we have validated that weIMPUTE incurs no additional computing costs compared to the imputation processing pipeline without the weIMPUTE framework.

## Overall design

weIMPUTE is a comprehensive imputation platform that streamlines the entire imputation process, from data preparation and phasing to imputing and post-quality control (QC) analysis (see Figure 1). By integrating these functionalities, weIMPUTE offers automated imputation without the need for additional data operations. The platform offers multiple pipelines to attend to various imputing scenarios, such as data segmentation and parallelization, while still allowing users to perform customized tasks, including phasing and imputing large datasets (Figure 2a-2c).

**Figure 1.**
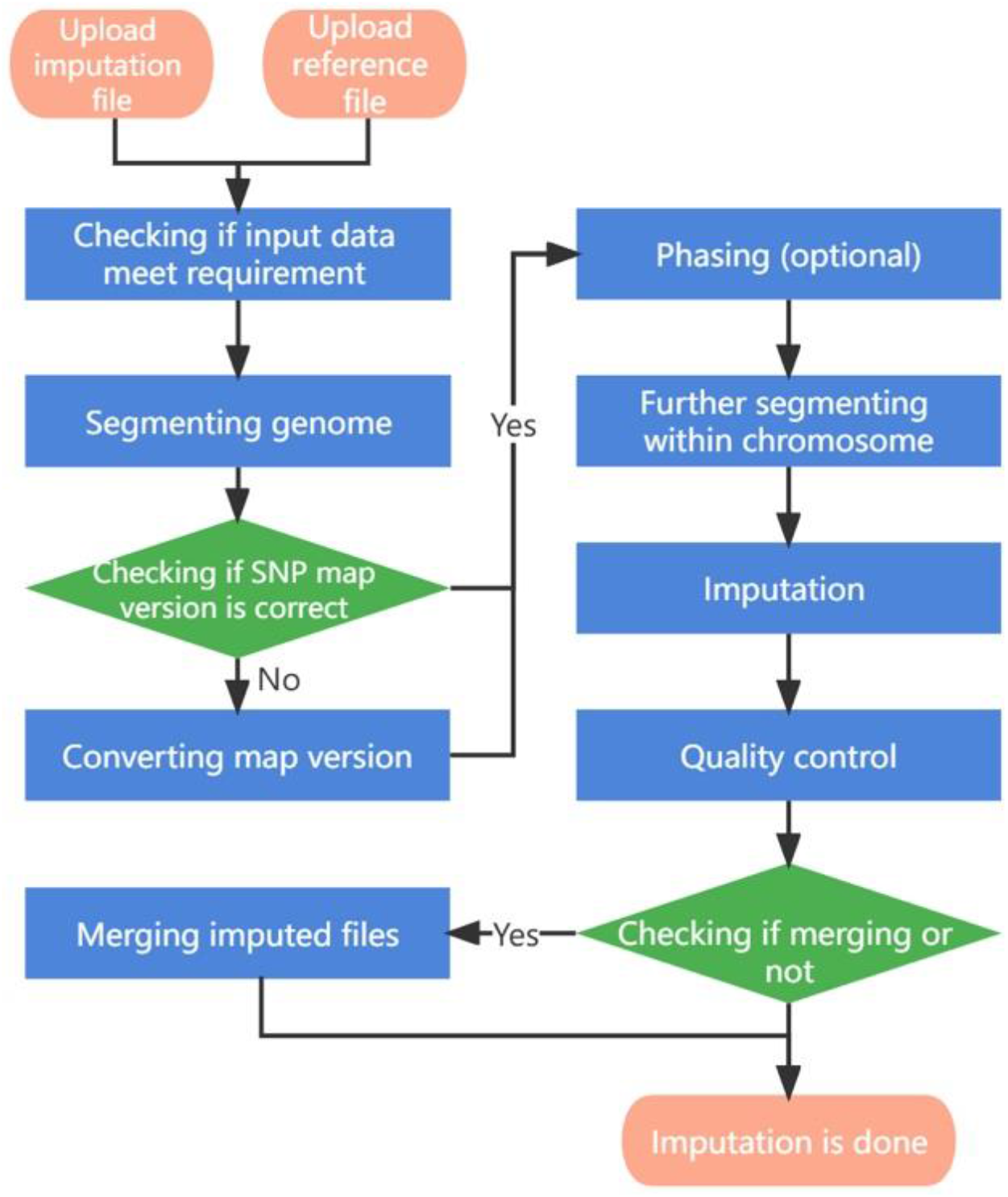
The workflow of phasing and imputation in weIMPUTE

**Figure 2.**
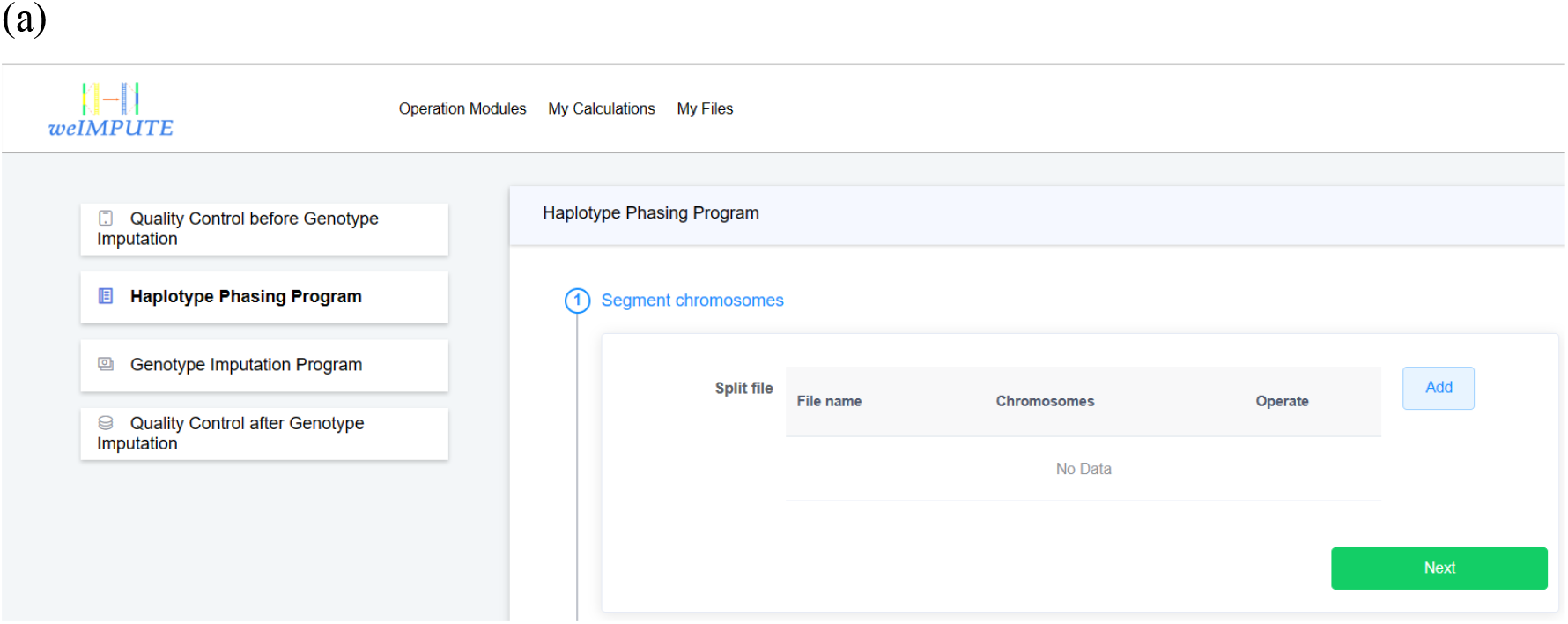
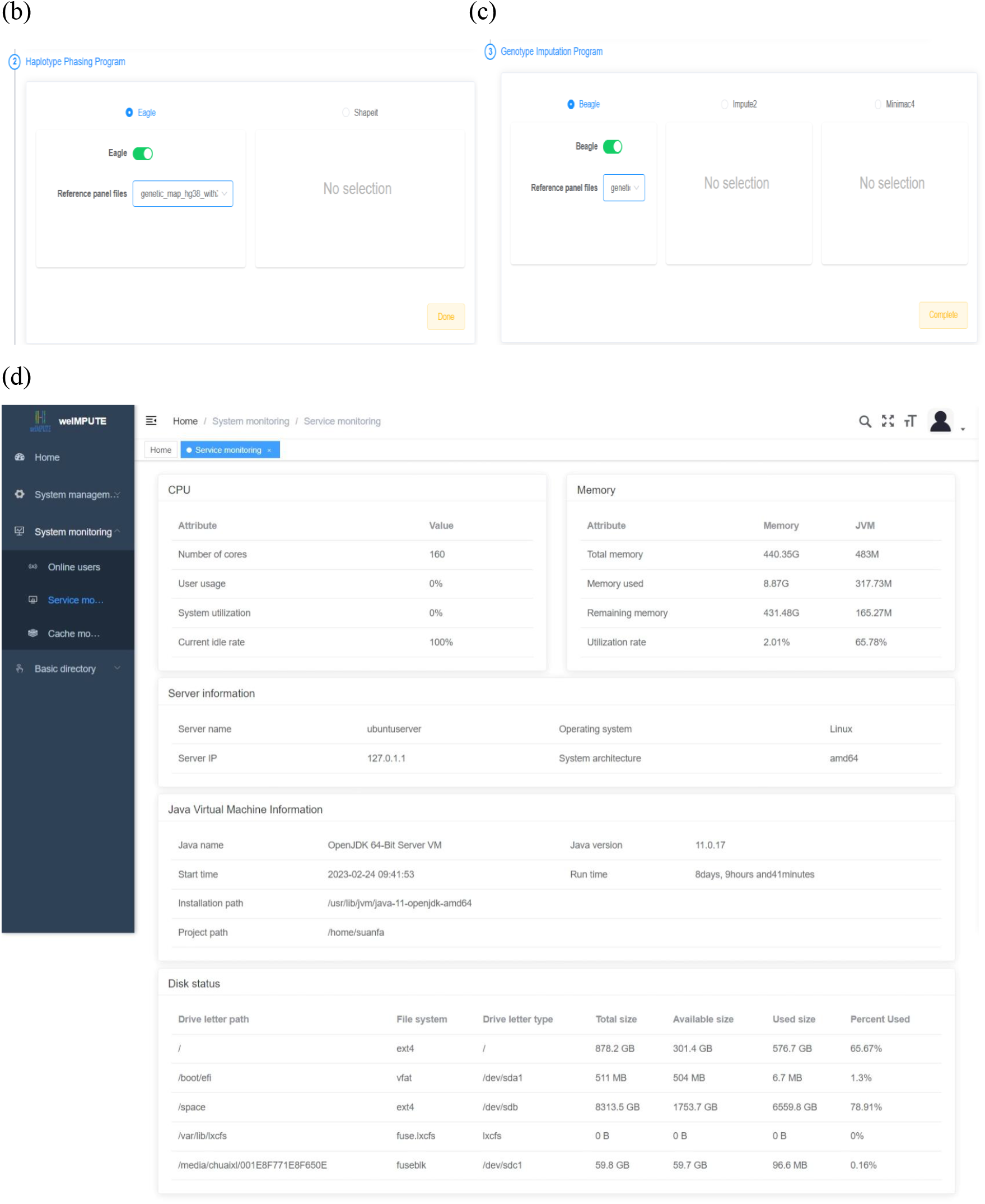
Performing data analysis with weIMPUTE. (a) weIMPUTE allows users to carry out customized tasks including quality control, phasing, and imputation with simple instructions and several clicks. (b) Options for phasing. Users could select Eagle2 or SHAPIT as their preferred phasing program. (c) Options for imputation. Users could choose Beagle5, IMPUTE2 or Minimac4 as their desired imputation program. (d) Management Module of weIMPUTE. weIMPUTE enables users to set up their personalized websites and to monitor and control service, cache, users, and data at backend.

Users can conveniently install weIMPUTE on their local machines and access it through a user-friendly website (Figure 2d), which also ensures data security through efficient datasets management. To ensure seamless compatibility across multiple operating systems (OS), all third-party libraries and software used in weIMPUTE have been packaged using Docker, eliminating the need for additional environment configuration. To achieve effective implementation, all modules within weIMPUTE are written in C++ and Java.

In the subsequent sections, we will delve into the key modules of the weIMPUTE pipeline, providing insights into its powerful and user-centric functionalities.

### Data Preparation

weIMPUTE accepts the widely used VCF format as input and output format for imputation. However, for Minimac4, which requires M3VCF format for the reference panel, VCF data will be converted to M3VCF using Minimac3 or m3vcftools. In cases where the human genome map versions of the reference panel and inference panel differ, the lift-over module performs the necessary conversion to align them. To comply with the phasing and imputing software requirements, the input data (inference panel) is automatically segmented into separate files by chromosome. For improved efficiency, users have the option to further segment data within each chromosome into smaller pieces (e.g., 5 million bp per file) using the segmentation module. If a segmented file contains too few SNPs, the QC module will issue a warning, prompting the user to remove it from the input file list. The QC module also verifies whether the input data meets the imputation requirements, such as being phased, map version consistency, and data format. If the input data includes sex chromosomes, they will be separated, and the pipeline for sex chromosome imputation will be executed automatically.

### Task Management

weIMPUTE offers a task management service that optimizes computing resources by default. Users are relieved of the burden of tuning numerous parameters for each phasing and imputation step, as the default settings are already optimized for efficient computing. Users have the freedom to select modules and determine their running order. Within each module, if multiple input files are present, they will be automatically assigned to different threads, optimized based on available computing resources. Task status can be monitored online or offline, and the storage management module handles automatic deletion of uploaded and output data files. Administrators can set a timer to determine when these files should be deleted.

### Imputed Data Merging and Filtering

After imputation, the post-QC module assesses the success of the imputation process. Users have the option to filter imputed variants based on imputation quality scores (r2 value). If desired, users can use the merge module to combine the filtered segment files into a single whole chromosome file.

### Platform Portability

weIMPUTE is containerized using Docker, making it cross-platform and easily accessible through a web browser on any OS that has Docker installed. For imputing large datasets, weIMPUTE can be installed on powerful computing nodes, allowing remote access through a laptop’s web browser. Users can check and modify the imputation job status at any time without the need to log into the computing node.

### Time Consumption and Memory Usage

Given that phasing and imputation are computationally intensive tasks, excessive computing resource usage can be a concern for GUI-based imputation software. To validate weIMPUTE’s effectiveness as an imputation service, we compared its time consumption and memory usage to traditional command line operations. The results show that weIMPUTE’s GUI-based approach exhibits similar time and memory costs as command line usage (Table 1 and 2). Leveraging Docker as a lightweight container, weIMPUTE effectively provides imputation services with optimal resource utilization.

**Table 1.** Summary of simulated data sets used in our experiments. Missing rate is the percentage of SNPs randomly masked to mimic random genotyping error.

| DataSet | Number of Samples | SNPs | Missing Rate (%) | VCF Size (MB) | VCF.gz Size (MB) |
| --- | --- | --- | --- | --- | --- |
| Reference Panel | 503 | 32515 | 0 | 63.73 | 4.02 |
| Data 1 | 450 | 32503 | 0.001 | 57.13 | 3.58 |
| Data 2 | 900 | 32503 | 0.001 | 113.35 | 6.42 |
| Data 3 | 1050 | 32503 | 0.001 | 131.96 | 7.23 |
| Data 4 | 1200 | 32503 | 0.001 | 150.56 | 8.1 |
| Data 5 | 1600 | 32503 | 0.001 | 200.18 | 10.6 |
| Data 6 | 2000 | 32503 | 0.001 | 249.79 | 13.16 |

**Table 2.**
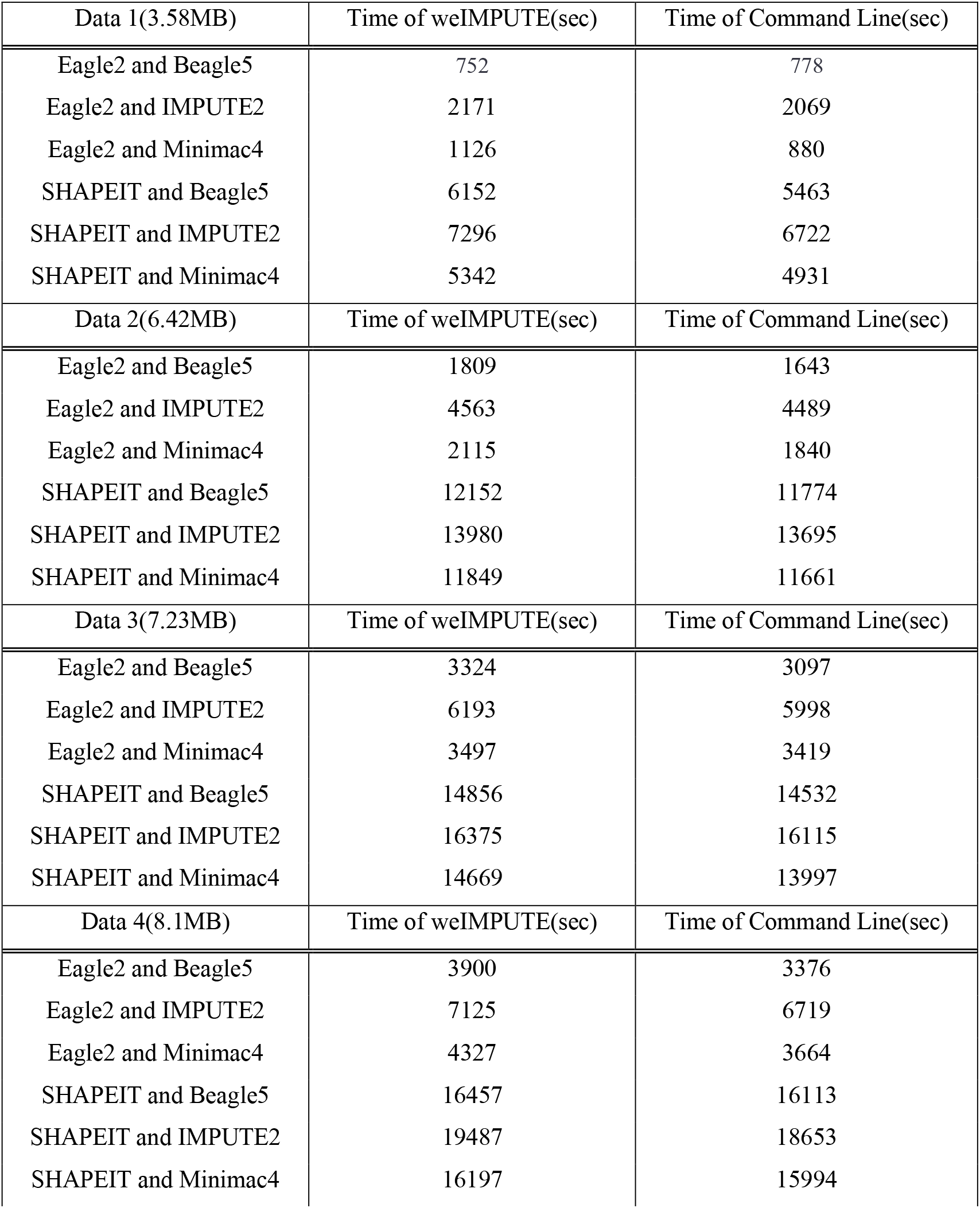
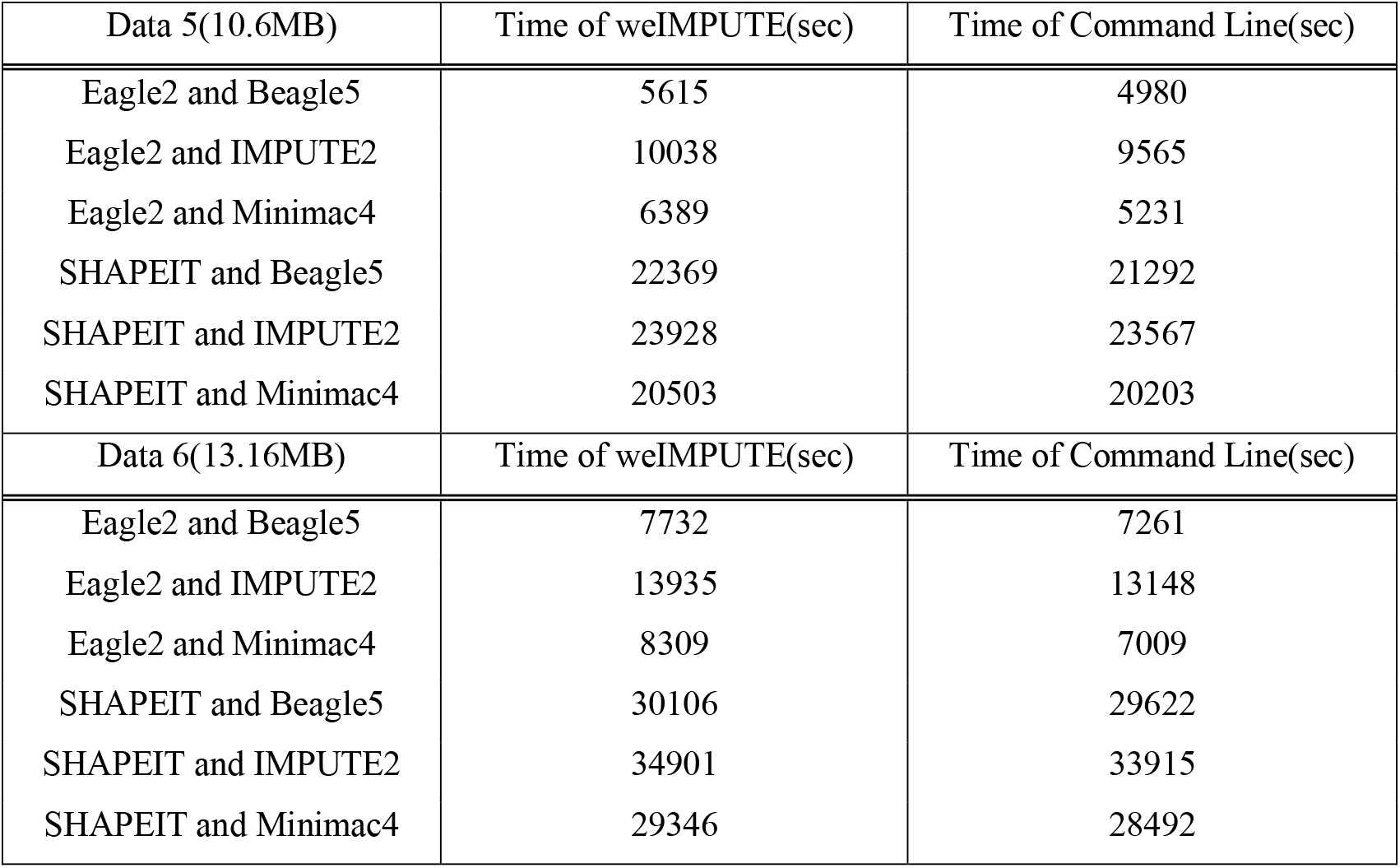
Phasing and imputation time comparison of weIMPUTE web service and command line version of various software combinations.

| Data 1(3.58MB) | Time of weIMPUTE(sec) | Time of Command Line(sec) |
| --- | --- | --- |
| Eagle2 and Beagle5 | 752 | 778 |
| Eagle2 and IMPUTE2 | 2171 | 2069 |
| Eagle2 and Minimac4 | 1126 | 880 |
| SHAPEIT and Beagle5 | 6152 | 5463 |
| SHAPEIT and IMPUTE2 | 7296 | 6722 |
| SHAPEIT and Minimac4 | 5342 | 4931 |
| Data 2(6.42MB) | Time of weIMPUTE(sec) | Time of Command Line(sec) |
| Eagle2 and Beagle5 | 1809 | 1643 |
| Eagle2 and IMPUTE2 | 4563 | 4489 |
| Eagle2 and Minimac4 | 2115 | 1840 |
| SHAPEIT and Beagle5 | 12152 | 11774 |
| SHAPEIT and IMPUTE2 | 13980 | 13695 |
| SHAPEIT and Minimac4 | 11849 | 11661 |
| Data 3(7.23MB) | Time of weIMPUTE(sec) | Time of Command Line(sec) |
| Eagle2 and Beagle5 | 3324 | 3097 |
| Eagle2 and IMPUTE2 | 6193 | 5998 |
| Eagle2 and Minimac4 | 3497 | 3419 |
| SHAPEIT and Beagle5 | 14856 | 14532 |
| SHAPEIT and IMPUTE2 | 16375 | 16115 |
| SHAPEIT and Minimac4 | 14669 | 13997 |
| Data 4(8.1MB) | Time of weIMPUTE(sec) | Time of Command Line(sec) |
| Eagle2 and Beagle5 | 3900 | 3376 |
| Eagle2 and IMPUTE2 | 7125 | 6719 |
| Eagle2 and Minimac4 | 4327 | 3664 |
| SHAPEIT and Beagle5 | 16457 | 16113 |
| SHAPEIT and IMPUTE2 | 19487 | 18653 |
| SHAPEIT and Minimac4 | 16197 | 15994 |

| Data 5(10.6MB) | Time of weIMPUTE(sec) | Time of Command Line(sec) |
| --- | --- | --- |
| Eagle2 and Beagle5 | 5615 | 4980 |
| Eagle2 and IMPUTE2 | 10038 | 9565 |
| Eagle2 and Minimac4 | 6389 | 5231 |
| SHAPEIT and Beagle5 | 22369 | 21292 |
| SHAPEIT and IMPUTE2 | 23928 | 23567 |
| SHAPEIT and Minimac4 | 20503 | 20203 |
| Data 6(13.16MB) | Time of weIMPUTE(sec) | Time of Command Line(sec) |
| Eagle2 and Beagle5 | 7732 | 7261 |
| Eagle2 and IMPUTE2 | 13935 | 13148 |
| Eagle2 and Minimac4 | 8309 | 7009 |
| SHAPEIT and Beagle5 | 30106 | 29622 |
| SHAPEIT and IMPUTE2 | 34901 | 33915 |
| SHAPEIT and Minimac4 | 29346 | 28492 |

## Conclusion

In summary, weIMPUTE offers an all-inclusive solution for imputation, encompassing data quality control and streamlining the entire imputation process. Its cross-platform compatibility and user-friendly interface make it a versatile tool, enabling imputation for data from diverse species. With weIMPUTE, researchers can seamlessly perform imputation without the need for extensive command line knowledge, and the platform’s automation ensures efficient and accurate results. This integration of functionalities and platform portability empowers researchers to harness the full potential of whole genome sequencing technology, expand sample sizes, and enhance statistical power in various research fields. weIMPUTE emerges as a valuable resource, advancing the field of genotype imputation and contributing to research advancements across different species and domains.

## Data Availability

The user manual, experiment and installation package of this study are available at: https://github.com/YOUTLab/weIMPUTE.

